# MIWI arginines orchestrate generation of functional pachytene piRNAs and spermiogenesis

**DOI:** 10.1101/2023.12.31.573779

**Authors:** Nicholas Vrettos, Jan Oppelt, Ansgar Zoch, Paraskevi Sgourdou, Haruka Yoshida, Brian Song, Ryan Fink, Dónal O’Carroll, Zissimos Mourelatos

## Abstract

N-terminal arginine (NTR) methylation is a conserved feature of PIWI proteins, which are central components of the PIWI-interacting RNA (piRNA) pathway. The significance and precise function of PIWI NTR methylation in mammals remains unknown. In mice, PIWI NTRs bind Tudor domain containing proteins (TDRDs) that have essential roles in piRNA biogenesis and the formation of the chromatoid body. Using mouse MIWI (PIWIL1) as paradigm, we demonstrate that the NTRs are essential for spermatogenesis through the regulation of transposons and gene expression. Surprisingly, the loss of TDRD5 and TDRKH interaction with MIWI results in defective piRNA amplification, rather than an expected failure of piRNA biogenesis. We find that piRNA amplification is necessary for both transposon control and for sustaining levels of select, nonconserved, pachytene piRNAs that target specific mRNAs required for spermatogenesis. Our findings support the notion that the vast majority of pachytene piRNAs are dispensable, acting as autonomous genetic elements that rely for propagation on MIWI piRNA amplification. MIWI-NTRs also mediate interactions with TDRD6 that are necessary for chromatoid body compaction. Furthermore, MIWI-NTRs promote stabilization of spermiogenic transcripts that drive nuclear compaction, which is essential for sperm formation. In summary, the NTRs underpin the diversification of MIWI protein function.

**Key points:**

- MIWI-NTRs coordinate interactions with TDRDs required for piRNA biogenesis to sustain piRNA amplification
- MIWI-NTRs are necessary for both transposon control and for sustaining levels of select pachytene piRNAs that target specific mRNAs required for spermiogenesis
- MIWI-NTRs mediate interactions with TDRD6 to compact the Chromatoid Body
- MIWI-NTRs underlie stabilization of spermiogenic transcripts that drive nuclear compaction, which is essential for sperm formation

## INTRODUCTION

PIWI proteins are expressed in the germline of all animals. They bind to PIWI-interacting RNAs (piRNAs), which guide them to complementary RNA targets (1). A cardinal function of piRNAs is to silence retrotransposons, thus protecting the genomic integrity of germ cells (2) (3) (4) (1). Mice express three Piwi proteins: MIWI (PIWIL1), MILI (PIWIL2) and MIWI2 (PIWIL4) (5) (6) (7). All three PIWI proteins are required for transposon silencing and male fertility (5) (6) (7) (8) (9) (10). Among the murine PIWI proteins, MILI and MIWI are piRNA-guided endonucleases (9) (11). In addition to transposon silencing, MIWI and pachytene piRNAs regulate gene expression by various mechanisms (5) (12) (13) (14) (15) (16) (17) (18) (19). Furthermore, MIWI loaded with pachytene piRNAs, TDRD6, and MVH, form the core of chromatoid body (CB) (20) (21) (22) (23) (12) (24). CBs first appears as small dispersed granules in secondary spermatocytes and coalesce into a single, large body (∼1 µm) in round spermatids (25) (26) (27) (20). CB is an aggresome (25) (23) and it is disposed as part of the cytoplasmic droplet during spermiogenesis (27) (28) (23) (12). Loss of either MIWI or TDRD6 leads to decompaction of CB (5) (20) (23) (12). In summary, PIWI proteins have acquired diverse genome protective and gene regulatory functions.

The N-terminus of PIWI proteins contains arginines (NTRs) that are symmetrically dimethylated (sDMA) by protein arginine methyltrasferase 5 (PRMT5) (29) and bind to Tudor domain containing proteins (TDRDs) (21) (30) (22) (31) (32). Multiple TDRDs interact with mouse PIWI proteins and have essential roles in piRNA biogenesis and function (33) (20) (34) (23) (35) (36) (37) (38) (39). TDRD1 interacts with MILI and is involved in selecting appropriate piRNA precursors for MILI piRNAs and MIWI2 loading (34). TDRKH (TDRD2) contains a transmembrane domain that tethers it to the cytoplasmic surface of mitochondria, and is essential for 3’ end trimming of MILI-bound prepachytene piRNAs (35) and for production of all MIWI-bound piRNAs (38). TDRKH interacts directly with MIWI, preferentially recognizing its unmethylated arginines (39). TDRD5 binds MIWI and piRNA precursors and is required for piRNA production from the entire length of the precursor transcripts (37). The sDMAs of MIWI are essential for mediating direct interactions to Tudor domains of TDRD6 (22). Deletion of TDRD6 leads to arrest at the elongation phase of haploid spermatids (step 12) (20) (23). TDRD6 does not have critical functions in piRNA biogenesis or piRNA-mediated transposon control but rather has a direct role in other CB-related functions of MIWI (23). How MIWI can coordinate such diverse functions remains unknown.

Here we report that MIWI-NTRs, by coordinating interactions with TDRDs required for piRNA biogenesis, orchestrate the functions of MIWI and piRNAs. Surprisingly, we find that the loss of TDRD5 and TDRKH interaction with MIWI results in defective piRNA amplification, rather than an expected failure of piRNA biogenesis. We demonstrate that MIWI piRNA amplification is necessary for both transposon control and for sustaining levels of select pachytene piRNAs that target specific mRNAs required for spermiogenesis. Our findings support the notion that the vast majority of pachytene piRNAs are dispensable, acting as autonomous genetic elements that rely for propagation on MIWI piRNA amplification. MIWI-NTRs also mediate interactions with TDRD6 that are necessary for CB compaction. Furthermore, MIWI-NTRs underlie stabilization of spermiogenic transcripts that drive nuclear compaction, which is essential for sperm formation. In summary, MIWI-NTRs enable functional diversification through the interaction with distinct TDRD proteins.

## RESULTS

### MIWI-NTRs are essential for fertility and spermiogenesis

To define the molecular function of MIWI’s NTR, we used genome editing of the endogenous *Miwi* locus to replace the six arginine residues (R4, R6, R8, R10, R12 and R14), which are subjected to symmetrical dimethylation, with lysines to generate the *Miwi^RK^* allele (**Fig. 1A** and **Supplementary Fig. 1A-D**). Unlike wild-type mice and heterozygous littermates, *Miwi^RK/RK^* males are infertile (**Fig. 1B, Supplementary Table 1)**, with 35% smaller testes (**Fig. 1C, Supplementary Table 2**) carrying no sperm in the seminiferous tubules or epididymis (**Fig. 1D**), due to a spermatid elongation failure phenotype and spermiogenesis arrest at step 8-9 (**Fig. 1E**). The MIWI^RK^ protein does not show any toxic or gain of function properties as it does not affect fertility when expressed along with wild-type MIWI in heterozygous *Miwi^+/RK^*males. The first wave of spermiogenesis in mice is synchronized, with round spermatids appearing at postnatal day 20 (P20) and spermatid elongation commencing around P26 (40) (41), allowing us to study MIWI^RK^ in viable germ cells prior to their arrest in *Miwi^RK/RK^*males. Unless otherwise indicated, all experiments were performed in P24 testes from heterozygous *Miwi^+/RK^* and homozygous *Miwi^RK/RK^* mice. To investigate whether the MIWI-RK mutation might have destabilized MIWI or other key piRNA protein factors, we performed Western Blots (WB) of testis lysates from *Miwi^+/RK^*and *Miwi^RK/RK^*. As shown in **Fig. 1F**, there were no changes in protein levels of MIWI or relevant piRNA factors in *Miwi^RK/RK^* mice.

**Figure 1.**
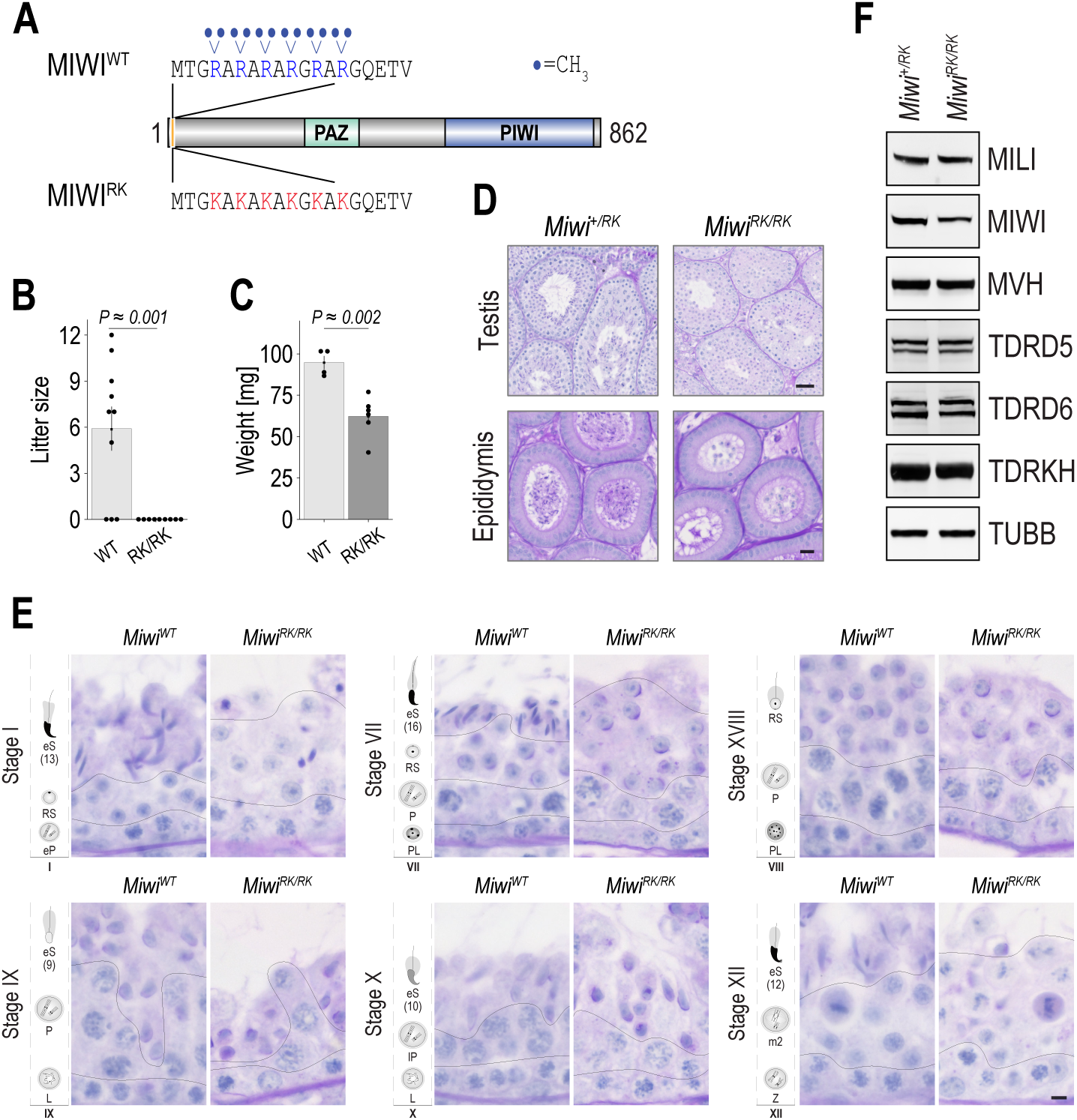
*Miwi^RK/RK^* mice are infertile due to spermatid elongation failure. **A.** Schematic representation of wild-type (WT) and arginine to lysine (RK) mouse MIWI proteins. **B.** Littersize per plug from *Miwi^WT^*and *Miwi^RK/RK^* studs mated to WT females and **C.** Adult testis weight; p-value of two-tailed Student’s t-test. **D.** Representative images of PAS-stained testis and epididymis sections from *Miwi^+/RK^* and *Miwi^RK/RK^*; scale bar, 20 µm. **E.** Spermatid elongation arrest at step 8-9. PAS-stained testis sections for different seminiferous cycle stages. PL, pre-leptotene; L, leptotene; Z, zygotene; eP, early pachytene; P, pachytene; lP, late pachytene; m2, secondary meiocyte; RS, round spermatid; eS (13), elongating spermatid (step 13). Scale bar, 5 µm. **F.** Western blots from indicated proteins in P24 testis lysates; beta-tubulin (TUBB) serves as loading control.

### MIWI-NTRs sustain amplification of pachytene cluster piRNAs and transposon control

To examine MIWI methylation, we probed MIWI immunoprecipitates (IP) from testes on WB with SYM11, an antibody that specifically recognizes sDMAs. As shown in **Fig. 2A**, SYM11 reactivity of MIWI is lost in *Miwi^RK/RK^*mice, consistent with R4, R6, R8, R10, R12 and R14 being the main arginines in MIWI that are subjected to symmetric dimethylation. Next, we examined with co-IP and WB assays whether interactions between MIWI-RK and TDRDs were altered. As shown in **Fig. 2B**, MIWI-RK fails to interact with TDRD5 and TDRKH.

**Figure 2.**
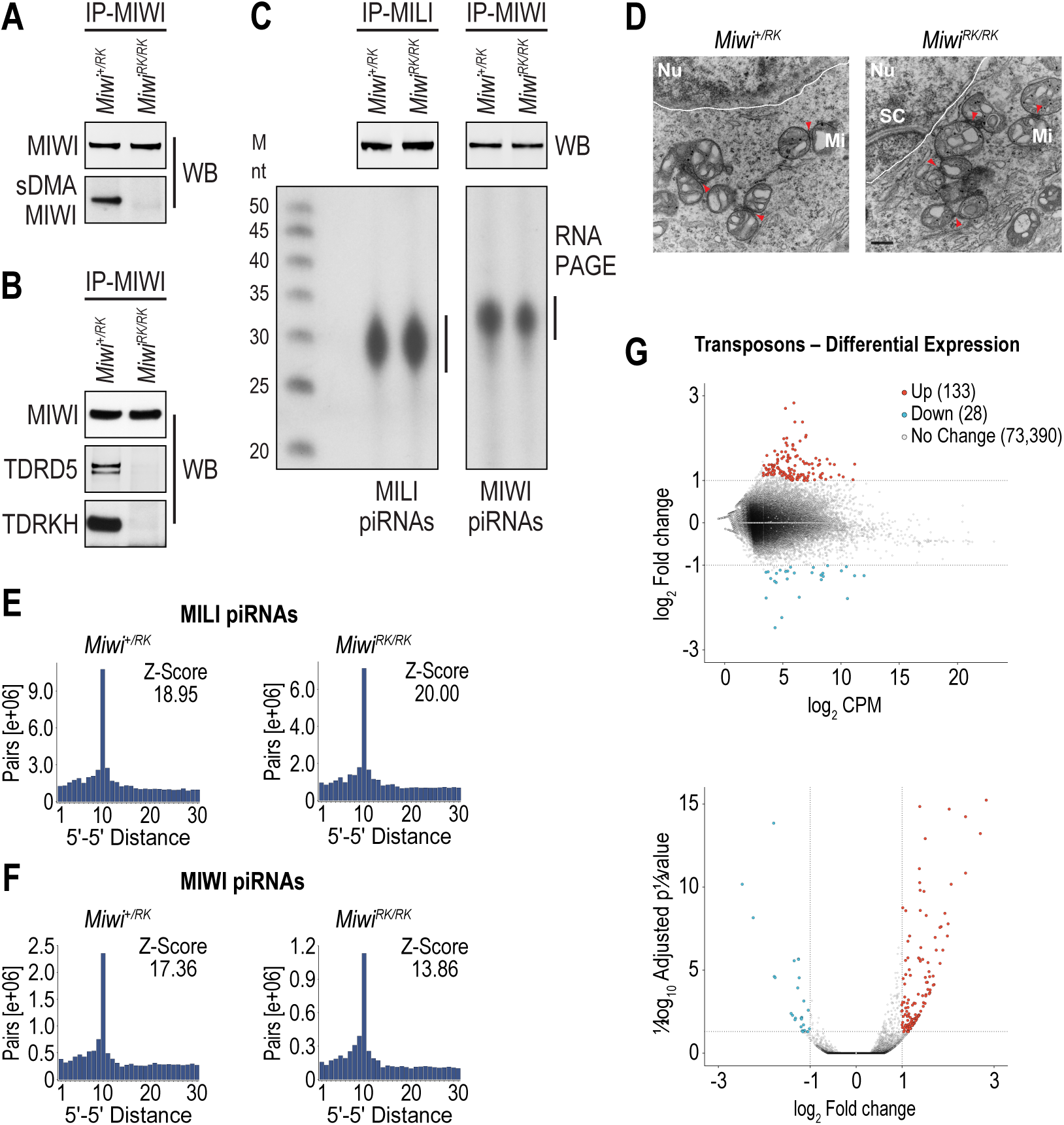
MIWI-NTRs sustain pachytene piRNA amplification of MIWI bound piRNAs and transposon control. **A, B.** Western blots of MIWI immunoprecipitates from indicated P24 testis lysates. **C.** Proteins (top) and bound piRNAs (bottom) of MILI and MIWI immunoprecipitates from indicated P24 testis lysates. **D.** Electron micrographs of *Miwi^+/RK^* and *Miwi^RK/RK^*pachytene spermatocytes; Nu, nucleus; dashed line, nuclear membrane; Mi, mitochondrion; red arrowheads, intermitochondrial cement; scale bar, 400 nm. **E, F.** 5’-5’ distance (ping-pong) analyses and Z-scores of MILI-bound (**E**) and MIWI-bound (**F**) piRNAs mapping to pachytene clusters in indicated genotypes. **G.** Differential expression analysis of transposons between *Miwi^+/RK^* and *Miwi^RK/^*^RK^ calculated from RNA-Seq libraries of P24 testes; top, MA plot (average expression relative to fold-change); bottom Volcano plot (statistical significance relative to fold-change); red, adjusted p-value < 0.05 and log_2_ fold-change >= 1; blue, adjusted p-value < 0.05 and log_2_ fold-change <=-1; grey, adjusted p-value >= 0.05 and/or log_2_ fold-change > −1 < 1.

Since MIWI-RK abolishes interactions with TDRDs that are essential for pachytene piRNA biogenesis, we hypothesized that piRNA populations in *Miwi^RK/RK^* mice would be affected. One possibility is that MIWI-RK would be devoid of most piRNAs since it does not interact with TDRKH, whose postnatal deletion leads to a near collapse of MIWI piRNAs (38). Another possibility is that the piRNA composition and genomic origin would be altered since MIWI-RK does not interact with TDRD5, which is required for piRNA generation from the entirety of pachytene piRNA precursors (37). We immunoprecipitated MILI and MIWI from *Miwi^+/RK^* and *Miwi^RK/RK^*lysates, isolated bound piRNAs and generated sequencing libraries (biological duplicates). Surprisingly, MIWI and MILI proteins are loaded with piRNAs in both genotypes (**Fig. 2C**), without any major differences in nucleotide composition (**Supplementary Fig. 2A**), piRNA sizes (**Supplementary Fig. 2B, C**), genomic origin (**Supplementary Figs. 3, 4, 5, 6**), or overall differential expression (**Supplementary Fig. 7**) indicating that MIWI-RK does not overtly affect piRNA biogenesis, precursor-transcript processing or piRNA loading. Ultrastructural analyses of spermatocytes in *Miwi^+/RK^*and *Miwi^RK/RK^*, shows that IMC is indistinguishable between the two genotypes (**Fig. 2D**). This is further indication that MIWI^RK^ does not have a major impact in piRNA biogenesis.

Next, we examined whether piRNA amplification (“ping-pong”) was affected by MIWI-RK by calculating the fraction of piRNAs in ping-pong pairs (**Supplementary Tables 3, 4, 5, 6, 7, 8**) and by plotting the 5’-5’ distance of MILI- and MIWI- bound piRNAs in *Miwi^+/RK^* and *Miwi^RK/RK^*. For MILI piRNAs, the 5’-5’ position analysis shows a peak at distance 10, with slightly increased Z-scores in *Miwi^RK/RK^*, consistent with largely intact MILI piRNA ping-pong (**Fig. 2E, Supplementary Fig. 8A**). In contrast, ping-pong Z-score values between MIWI bound piRNAs are reduced in *Miwi^RK/RK^*, indicative of attenuated piRNA amplification by MIWI^RK^ (Z-score of 13.86 in *Miwi^RK/RK^* versus 17.36 in *Miwi^+/RK^* for piRNAs mapping to pachytene clusters **Fig. 2F**; Z-score of 11.63 in *Miwi^RK/RK^* versus 13.26 in *Miwi^+/RK^* for piRNAs mapping to gene exons in antisense orientation; Z-score of 13.15 in *Miwi^RK/RK^* versus 16.19 in *Miwi^+/RK^* for piRNAs mapping to gene exons in sense orientation, **Supplementary Fig. 8B**). Z-scores were very similar for heterotypic, MILI-MIWI ping-pong between the two genotypes (**Supplementary Fig. 9**). To examine the impact of attenuated ping-pong in transposons, we analyzed the levels of individual transposable elements (TEs) by performing RNA-Seq from total RNA isolated from *Miwi^+/RK^* and *Miwi^RK/RK^*testes. While the levels of the vast majority of TEs are not altered between the two genotypes, we find that five times as many TEs are upregulated than downregulated in *Miwi^RK/RK^* (133 upregulated, 28 downregulated, **Fig. 2G** and **Supplementary Table 9**, adj. p-value < 0.05, fold change >=2), indicating that attenuated piRNA amplification decreases clearance of select TEs in *Miwi^RK/RK^* mice.

### MIWI^RK^ impacts the mRNA transcriptome but not the translatome

To investigate the possible impact of MIWI^RK^ on the expression and translation of mRNAs, we performed, in biological triplicates, ribosome profiling (Ribo-Seq) from *Miwi^+/RK^* and *Miwi^RK/RK^*testes and analyzed the libraries along with concurrent RNA-Seq. Although the expression levels of most transcripts are not altered between the two genotypes, we find that 180 (116 protein-coding) genes are upregulated and 71 (38 protein-coding) genes are downregulated in *Miwi^RK/RK^* (**Fig. 3A** and **Supplementary Table 10**; adj. p-value < 0.05, fold change >=2). We find that the ribosome occupancy is essentially identical in both genotypes, with changes in only a handful of genes. Specifically, 14 transcripts (10 genes) were more occupied and 15 (12 genes) less occupied (adj. p-value <0.05, fold change >=2), indicating that mRNA translation is not grossly impacted in *Miwi^RK/RK^* mice (**Fig 3B** and **Supplementary Table 11**).

**Figure 3.**
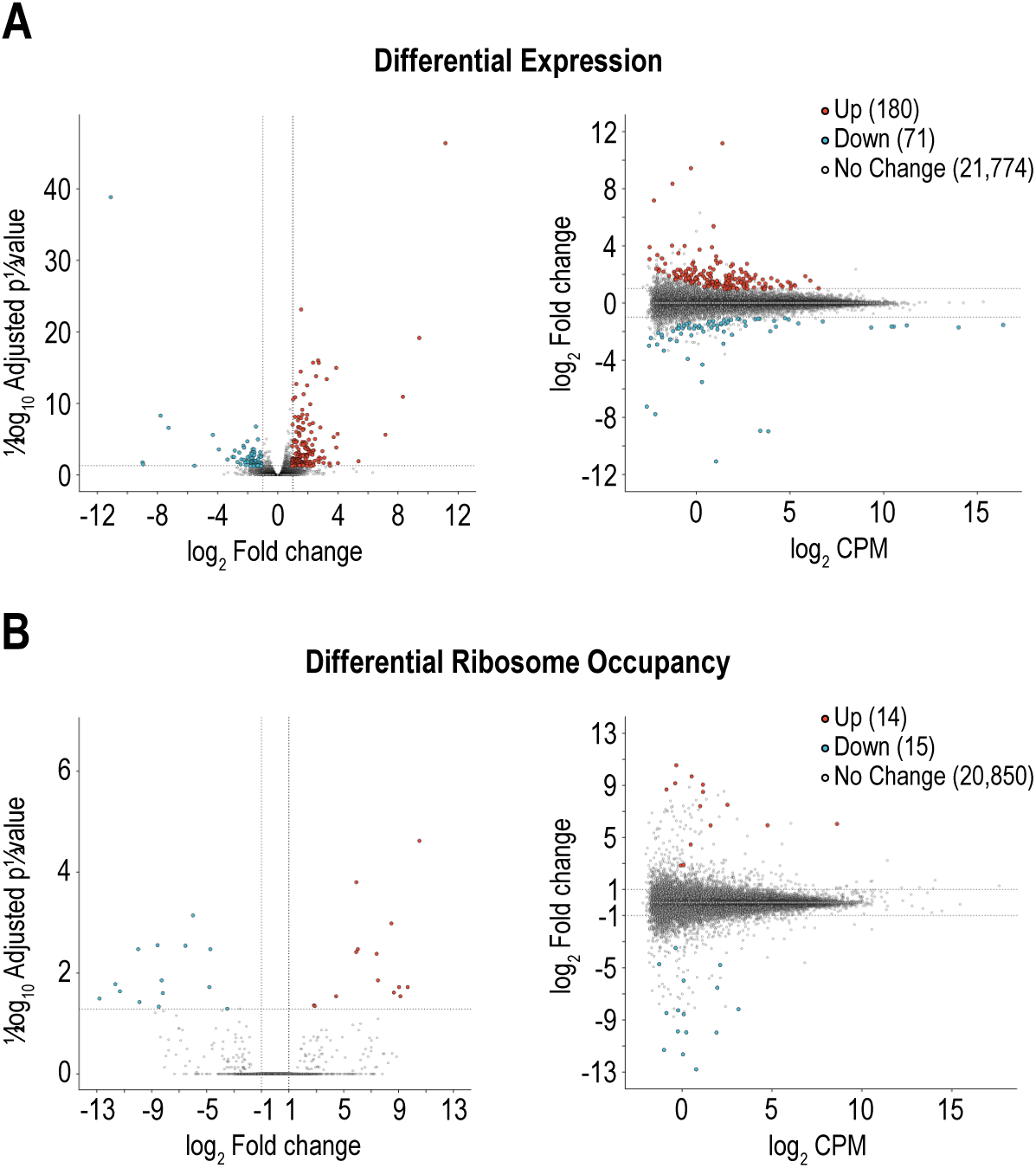
MIWI^RK^ impacts the transcriptome but not the translatome. **A, B.** Differential expression analysis calculated from RNA-Seq (**A**) and differential ribosome occupancy calculated from Ribo-seq (**B**) between *Miwi^+/RK^* and *Miwi^RK/RK^* P24 testes, visualized by Volcano (left) and MA (right) plots; red, adjusted p-value < 0.05 and log_2_ fold-change >= 1; blue, adjusted p-value < 0.05 and log_2_ fold-change <=-1; grey, adjusted p-value >= 0.05 and/or log_2_ fold-change > −1 < 1.

### MIWI-NTRs sustain piRNAs that cleave and destabilize select mRNAs essential for spermatogenesis

Since piRNA amplification is attenuated by MIWI-RK, we hypothesized that some of the upregulated transcripts in *Miwi^RK/RK^*could be directly targeted and cleaved by piRNAs with decreased levels in the mutant. To investigate our hypothesis, we first predicted RNA targets that are potentially cleaved by piRNAs. We used parameters that considered the requirement for both extended stretches of complementarity between the piRNA and its target, and tolerance for mismatches (9) (42) (43) (44). We find that of the 45,029 transcripts expressed at P24 testes, 31,745 (∼70%) are theoretically targeted by 46,494 out of 226,198 unique MIWI-bound piRNAs. The meta-transcript distribution of predicted target sites in meta-mRNA is heavily skewed towards the 3’ Untranslated Region (3’-UTR, **Fig. 4A**).

**Figure 4.**
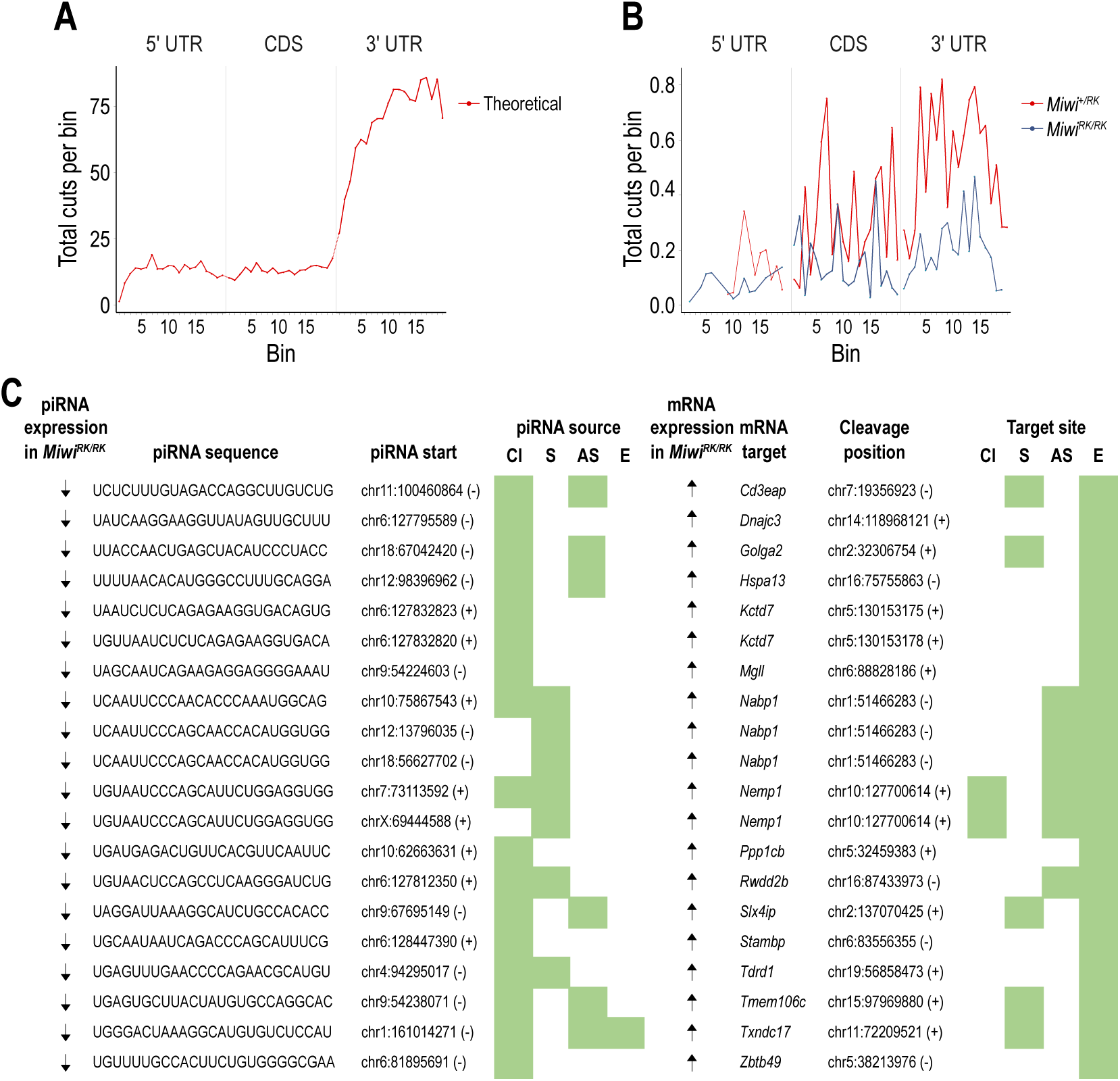
MIWI-NTRs sustain piRNAs that cleave and destabilize select mRNAs essential for spermiogenesis. Metatranscript distribution of predicted, piRNA-mediated cleavages on mRNA targets, without (“theoretical”, **A**), or with degradome-Seq support (**B**). **C.** Sequence and genomic location of piRNAs found to cleave and destabilize the listed mRNAs at the indicated coordinates. piRNAs and cleaved RNA fragments map to the following categories (green boxes): Cl: piRNA cluster; S: sense-aligning repeat sequence; AS: antisense-aligning repeat sequence; E: exonic sequence. Arrows indicate changes in the expression of piRNAs (down) and mRNA targets (up) in *Miwi^RK/RK^* compared to *Miwi^+/RK^*.

PIWI nucleases cleave the phosphodiester bond of target RNAs across from the 10^th^ and 11^th^ nucleotide of their bound piRNA, generating a decay fragment that has a 5’ phosphate group, which can be captured and sequenced with Degradome-Seq. We performed Degradome-Seq from poly(A) RNA isolated from *Miwi^+/RK^* and *Miwi^RK/RK^*testes (biological duplicates). To identify which of the predicted target sites were cleaved by piRNAs, we analyzed the position of the first nucleotide of decay intermediates captured with Degradome-Seq in relationship with piRNA-target RNA pairs. We find that the targeting of 686 transcripts (678 genes out of 31,745; 2%) by 662 unique, MIWI-bound piRNAs, is supported by Degradome-Seq (meta-transcript distribution of target sites with Degradome-Seq support in the two genotypes is shown in **Fig. 4B**). More than half (374 out of 662) of the unique MIWI-bound piRNAs, originate from repeat-related loci in either sense or antisense orientation to the repeat annotation. The predicted piRNA target sites are primarily in 3’ UTRs (764 target sites), while Coding Sequence (CDS) and 5’ Untranslated Region (5’ UTR) are much less favored (170 and 29 target sites, respectively).

Next, we identified MIWI-bound piRNAs whose expression is lower in *Miwi^RK/RK^*, compared to *Miwi^+/RK^*, and with corresponding mRNA targets upregulated. We find that of 678 genes targeted by piRNAs, 17 are upregulated (adj. p-value <0.1, fold change >0) in *Miwi^RK/RK^*with corresponding downregulation (adj. p-value <0.25, fold change >0) of 19 targeting, MIWI-bound piRNAs (**Fig. 4C**). Notably, the sequences of half of the piRNA-target RNA pairs are derived from repeat elements while few piRNAs targeted piRNA cluster transcripts (**Fig. 4C**). Strikingly, 3 of the 17 upregulated genes in *Miwi^RK/RK^* along with the corresponding downregulated targeting piRNAs, are the same as those identified in the *pi6* (*Dnajc3*, *Kctd7*) and *pi18* (*Golga2*) knockouts (18) (19) (**Fig. 4C**). Another two (*Stambp2* and *Tdrd1*) were previously shown to be upregulated in *Miwi-*null mice (13). GOLGA2 (GM130) is a Golgi matrix protein essential for acrosome formation (45). Ultrastructural examination of *Miwi^+/RK^* and *Miwi^RK/RK^* testes shows that the acrosome is intact in *Miwi^+/RK^* spermatids, with normal formation of acroplaxome, acrosomal granule and acrosomal vesicle (**Supplementary Fig. 10**). However, acrosome is not formed in *Miwi^RK/RK^* spermatids, which in most cases show a mis-oriented Golgi apparatus that does not form an acrosome or less frequently multiple, small fragmented acrosomal granules that do not coalesce to form an acrosome (**Supplementary Fig. 10**). The phenotype of *Miwi^RK/RK^* is more similar to that of *Miwi*-null (5) (12) (13) and much more severe than that seen with loss of individual piRNA loci (18) (19), consistent with the larger transcriptome alterations found in *Miwi^RK/RK^* mice.

### MIWI-NTRs are required for TDRD6 interaction, chromatoid body compaction and the stability of key spermiogenic transcripts

Next, we tested with co-IP and WB assays whether the binding between MIWI-RK and TDRD6 was abolished. As shown in **Fig. 5A**, MIWI-RK fails to interact with TDRD6. We examined by immunofluorescence (IF) the localization of MIWI and TDRD6 in testes from heterozygous and homozygous *Miwi^RK^* mice. As shown in **Fig. 5B**, there is similar localization and CB formation, in both genotypes. However, ultrastructural analyses show that CB is less compact, less electron dense and often fragmented in *Miwi^RK/RK^*round spermatids (**Fig. 5C**), indicating that MIWI-TDRD6 interaction is required for CB compaction. The CB defect in *Miwi^RK/RK^*is similar to what is seen in deletion mutants of *Miwi* or *Tdrd6* (5) (20) (23).

**Figure 5.**
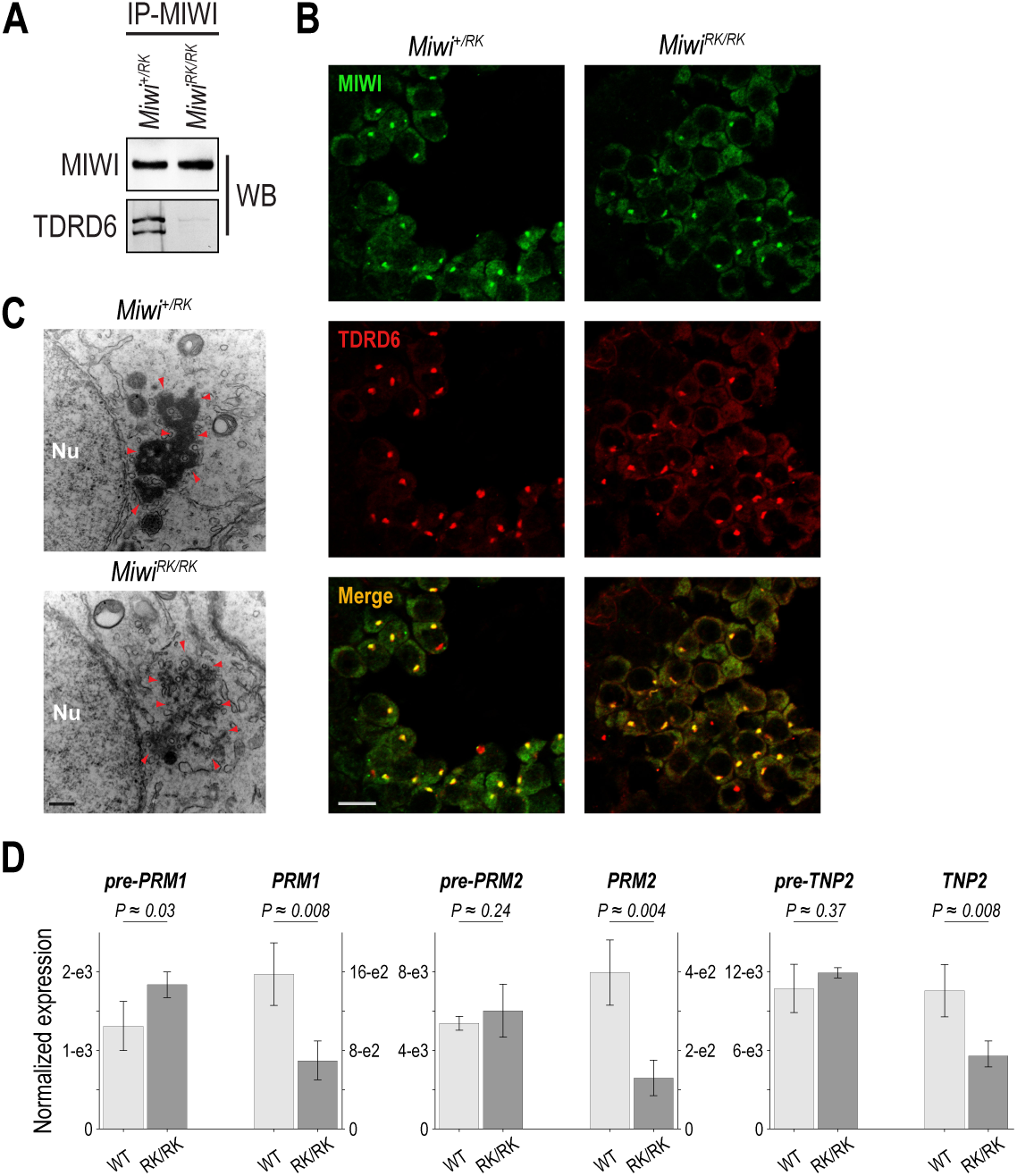
MIWI-NTRs are required for TDRD6 interaction, chromatoid body compaction and stability of key spermiogenic mRNAs. **A.** Western blots of MIWI immunoprecipitates from indicated P24 testis lysates. **B.** Immunofluorescence staining for MIWI (green) and TDRD6 (red) on round spermatid populations detected in mouse sections of *Miwi^+/RK^*and *Miwi^RK/RK^* P24 testis sections. Scale bar, 10 µm. **C.** Electron micrographs of *Miwi^+/RK^*and *Miwi^RK/RK^* round spermatids; Nu, nucleus; red arrowheads, chromatoid body; scale bar, 400 nm. **D.** pre-mRNA and mRNA relative expression levels for indicated genes, assayed by RT-qPCR (biological and technical triplicates) in *Miwi^+/RK^* and *Miwi^RK/RK^* P24 testes. Assays for *Tcp1* were used to normalize for loading and differentiation stage; p-values of one-sided t-test.

Among the 38 downregulated genes in *Miwi^RK/RK^* (**Supplementary Table 10**) many are essential spermiogenic genes that were previously shown to be downregulated in *Miwi*-null mice (5) (12). These include genes coding for nuclear transition proteins (*Tnp2)* and protamines (*Prm1*, *Prm2*), which replace the histones during spermatid elongation to form the sperm nucleus (46) (41). To examine whether TDRD6 has similar impact on the transcriptome, we analyzed previously published RNA-Seq data from sorted P26 spermatids from *Tdrd6 ^−/+^* and *Tdrd6 ^−/–^* mice (47) and compared them to *Miwi^RK/RK^*. We find 6 downregulated genes (adj. p-value <0.05, fold change >=2) that are common to both *Tdrd6*-null and *Miwi^RK/RK^* mice, including *Tnp2*, *Prm1* and *Prm2* (**Supplementary Table 12**). To examine whether the reduced transcript levels in *Miwi^RK/RK^* were secondary to reduced production or destabilization, we performed RT-qPCR, in biological and technical triplicates, and utilized the *Tcp1* transcript for input and differentiation stage normalization (48). We find that mRNA levels for *Prm1*, *Prm2* and *Tnp2* are significantly reduced in *Miwi^RK/RK^* compared to in *Miwi^+/RK^*, while their pre-mRNA levels are either unchanged or slightly upregulated **( Fig. 5D)**, indicating mRNA destabilization of key spermiogenic transcripts by MIWI-RK. Collectively, these findings demonstrate that downregulation of spermiogenic transcripts required for chromatin remodeling and sperm nuclear compaction is a shared molecular phenotype of mice null for either *Miwi* or *Tdrd6*, as well as *Miwi^RK/RK^*mice, which are unable to form MIWI-TDRD6 complexes.

## DISCUSSION

Prior genetic investigations of MIWI and TDRDs relied on gene knockouts and illuminated key aspects of piRNA biology but could not directly address the role of MIWI-TDRD interactions in piRNA biogenesis and function, *in vivo*. Here, with the separation-of-function *Miwi^RK^* allele we specifically uncouple MIWI from TDRDs, while preserving expression of all TDRDs and functions associated with all other PIWI domains. A proposed model that emerges from our findings is shown in **Figure 6** and discussed below.

**Figure 6.**
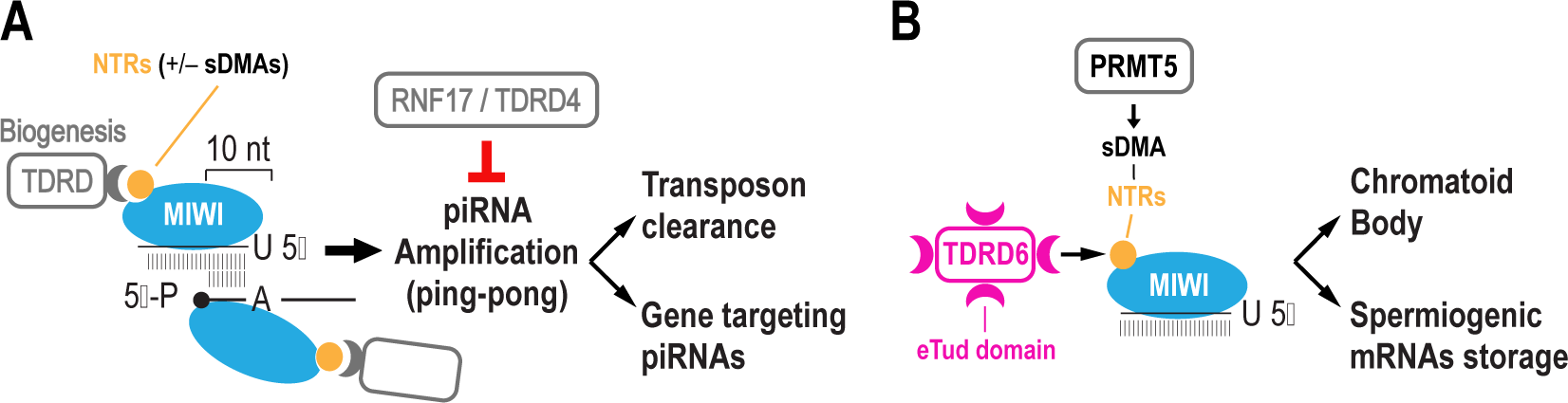
Model of MIWI-NTR functions. **A, B.** MIWI-NTR function in piRNA amplification, transposon clearance and gene targeting (**A**), and in chromatoid body compaction and spermiogenic mRNA stability (**B**).

We find that MIWI-RK does not impact primary piRNA biogenesis but attenuates piRNA amplification, leading to reduced transposon clearance. By disrupting the interaction of MIWI with TDRDs that are involved in piRNA biogenesis, we unveil the few functional piRNAs that target specific mRNAs for destabilization. Amongst them are previously reported gene targets (13) and piRNAs from the *pi6* and *pi18* loci, recently shown to be essential for mouse fertility (18) (19). We also uncover targeting but “nonfunctional” piRNAs that appear to cleave mRNAs without impacting their stability or translation. We find that the sequences of most piRNAs that cleave mRNAs (both functional and “nonfunctional”) are related to repeat elements. In turn, these piRNAs target repeat element sequences embedded in mRNAs, typically in their 3’-UTRs. It is not clear why some mRNAs cleaved by piRNAs are not destabilized. We favor the idea that such cleavages are infrequent events that affect only a small fraction of mRNA molecules. This could be due to a number of factors, such as low levels of the targeting piRNA and/or large number of target sites distributed amongst TE transcripts and the targeted mRNAs, diluting the impact of targeting piRNAs on the protein-coding transcriptome. We find that the vast majority of pachytene piRNAs are not involved in gene regulation, consistent with our previous hypothesis (12) and with more recent reports (49) (18) (19).

Our findings support the primacy of MIWI piRNA amplification for postmeiotic clearance of TE transcripts (9). They also support the notion that the vast majority of pachytene piRNAs act as autonomous genetic elements that rely for propagation on few functional piRNAs (18) (19). We propose that production of these functional piRNAs depends on MIWI piRNA amplification, which in turn is regulated by MIWI-NTRs. Since piRNA amplification involves mostly repeat-related elements whose sequences are divergent in different mammalian species, the functional piRNAs that arise are not conserved. Our proposition is further supported by the demonstration that RNF17/TDRD4 (33) is a negative regulator of meiotic piRNA amplification (36). In the absence of RNF17, hyperactive piRNA amplification leads to inappropriate targeting and downregulation of multiple protein-coding genes (36) and spermiogenic arrest at the round spermatid stage (33) (36) (**Figure 6A**).

We have previously shown that MIWI is a component of spermiogenic mRNPs that stabilize stored mRNAs, notably *Tnp2, Prm1*, *Prm2* whose translation is delayed until spermatid elongation (12). By analyzing published RNA-Seq data from sorted P26 spermatids of *Tdrd6^−/+^* and *Tdrd6^−/–^* mice (47), we find that the same genes are downregulated in the absence of TDRD6. We show that the interaction of MIWI with TDRD6 is required for CB compaction. Furthermore, we solidify the role of MIWI and its NTRs in stabilization of these key spermiogenic mRNAs that drive histone replacement of spermatid nuclei. The molecular mechanism that underlies spermiogenic mRNA stabilization remains to be determined but likely involves a non-cleaving, mRNA binding mode for MIWI. We previously reported that MIWI binds directly to translationally repressed spermiogenic mRNAs (by CLIP-Seq and CIMS –crosslinked induced mutation site– analyses) and proposed that it does so without using piRNAs (12). It is also possible that MIWI piRNAs bind spermiogenic mRNAs with partial complementarity that does not lead to mRNA cleavage. We have shown that such a binding mode occurs in *Drosophila melanogaster*, where Aub-piRNAs bind mRNAs with partial complementarity in the germ plasm and along with Tudor protein form germ granules (50). Further studies are required to identify the precise molecular function of MIWI in spermiogenic mRNP assembly and stabilization.

Our findings draw parallels between the functions of mouse MIWI/TDRDs and *D. melanogaster* Aub/Tdrds. We recently examined, genetically, the role of Aub-NTRs and their methylation status in piRNA biogenesis, amplification and formation of germ granules (51). The latter consist of mRNPs that contain mRNAs, bound directly by Aub (50), and also contain Tudor (50) (52) (53), Vasa (54) (55) (56) and other RNA-binding Proteins (57). Germ granules assemble at the posterior of *D. melanogaster* oocytes and their mRNAs are essential for primordial germ cell specification in the developing embryos (57). By genetic deletion or germ cell specific knockdown of *Drosophila Prmt5*, we found that germ granules never formed as Aub with unmethylated NTRs could not interact with Tudor (51). piRNA biogenesis, amplification and transposon control were intact in flies expressing endogenous Aub with unmethylated NTRs (51). However, by mutating Aub-NTRs to lysines, we found that germ granule formation, piRNA amplification and transposon control all collapsed, as the Aub-RK mutant protein was unable to interact not only with Tudor but also with Tdrds involved in piRNA amplification (51). Primary piRNA biogenesis was not impaired by Aub-RK (51). We think that similar functional interactions take place in mice between MIWI and TDRDs. MIWI-RK, by disrupting interactions with TDRDs involved in piRNA biogenesis and with TDRD6, leads to attenuation of piRNA amplification, TE transcript upregulation, CB decompaction and spermiogenic mRNA destabilization. We speculate that sDMAs of MIWI-NTRs and the valency of their interactions with the multiple eTUD domains of TDRD6, drive functions relating to CB compaction and spermiogenic mRNP formation.

While our manuscript was under revision, Wei et al. reported the generation of a MIWI-RK mouse mutant, with spermiogenesis arrest at step 8 (58), similar to what we find. Wei et al. characterized their allele in a *Miwi null* background (*Miwi^RK/–^*) and found that MIWI-RK protein is reduced with concomitant reduction of their bound pachytene piRNAs, which are qualitatively similar to those bound by wild-type MIWI (58). We also find that MIWI-RK does not overtly affect pachytene piRNA biogenesis or piRNA loading but we do not observe the large reduction in the levels of MIWI-RK protein or piRNAs, since we characterize mice that are homozygous for the mutant allele (*Miwi^RK/RK^*). Importantly, by performing deep characterization of our MIWI-RK mice, with RNA-Seq, Ribo-Seq, Degradome-Seq, piRNA-Seq and extensive bioinformatical analyses, we provide molecular explanations for the role of MIWI NTRs in spermiogenesis and pachytene piRNA function.

In summary, we find that a central molecular mechanism by which PIWI proteins are directed and utilized by distinct cellular pathways, resides in the NTR domain. This mechanism is essential for the integrity and continuity of the germline.

## MATERIALS AND METHODS

### Generation and genotyping of the *Miwi^RK^* allele

The mouse *Miwi^RK^* allele was generated through CRISPR-CAS9 gene editing on fertilized 1-cell stage zygotes of B6CBAF1/Crl genetic background as previously described (59),(60). *Cas9* mRNA was injected together with a small guide RNA (CTGACCTCGTGCCCTGCCGC) targeting the MIWI-NTR encoding exon plus a 200 nt ssDNA donor oligonucleotide (GTGCTTTAAAAAGGTTTAGTGATAAAATGGTGAATGCACGTGAGCCCATCGTGTGCTTTTCCTCTCTGACAGAAAATGACTGGCaagGCCaagGCTaaaGCCaagGGCaaaGCAaagGGTCAGGAGACGGTGCAGCATGTTGGGGCTGCTGCGGTGAGTACCATTCTTATATAGCTCAATAGCATCTTAACAACCAGCCA) delivering the expected nucleotide changes flanked by two 5’ and 3’ homology arms, 84 nt and 83 nt in length, respectively **(Supplementary Figure 1A, B**). F_0_ offspring carrying the *Miwi^RK^* allele were identified by PCR and validated with Sanger sequencing (**Supplementary Figure 1C, D**). *Miwi^RK^*line was established from one F_0_ founder mouse and was back-crossed several times onto a C57BL6/N genetic background. Therefore, transgenic mice in this study, were maintained on a mixed B6CBAF1/Crl; C57BL6/N genetic background.

Mouse genotyping was determined with PCR (90 s at 95°C, followed by 33 cycles of 15 s 95°C, 30 s 60°C and 50 s 72°C) using the following primers: outer-fw (CTGGAACTGATGGTTTCGTGCC), outer-rv (CTAGTCCACAGCATTCAGGGACTG), wt-fw (TGACAGAAAATGACTGGCCGA) and mut-rv (CCTTTGCTTTGCCCTTGGCT). A common amplicon is produced (743 bp) as well as, a *Miwi^WT^*-(345 bp) and/or a *Miwi^RK^*-specific (450 bp) (**Supplementary Figure 1D**).

### Mouse experimentation

Male fertility was assayed by timed matings of an adult stud male with a wild-type C57BL6/N adult female for 4 d and separation of the female upon detection of a vaginal plug. The plug date was counted as embryonic day 0.5 (E0.5) and the number of embryos at E16.5 was counted for each plugged female. Animal tissue samples were collected from one or more litters per experiment and allocated to groups according to genotype. No further randomization or blinding was applied during data acquisition and analysis. Animals were maintained at the University of Edinburgh, UK, in accordance with the regulation of the UK Home Office. Ethical approval for the mouse experimentation has been given by the University of Edinburgh’s Animal Welfare and Ethical Review Body and the work was done under license from the United Kingdom’s Home Office. Furthermore, mouse experimentation for this project was reviewed and approved by the Institutional Animal Care and Use Committee (IACUC) of the University of Pennsylvania.

### Histology and immunohistochemistry

For periodic acid Schiff (PAS) staining, testes and epididymis were fixed overnight in Bouin’s fixative (Millipore Sigma) followed by several washes in 70 % ethanol and embedding in paraffin. Next, 3 µm sections were cut on a microtome (Leica), rehydrated through an alcohol series according to standard laboratory procedures and then stained with a PAS staining kit (TCS Biosciences) according to the manufacturer’s instructions. Stained sections were rehydrated through a reverse alcohol series and mounted on coverslips with Pertex mounting medium (Pioneer Research Chemicals) according to standard laboratory procedures. Slides were imaged on Zeiss AxioScan scanning microscope at 40× magnification. Cropped images were exported using the Zeiss Zen software and further processed in ImageJ.

For IF, testes were fixed overnight in 4% PFA and embedded in paraffin. The testicular tissue was cut into 5 µm thick sections on a microtome (Leica), mounted onto Superfrost Plus microscope slides (Fisherbrand) and left to dry overnight at 47°C. The tissue was dewaxed by immersion in xylene three times for 5 min each, rehydrated sequentially in 100% ethanol and finally rinsed in distilled water (dH_2_O). Slides were placed in Coplin jars filled with antigen retrieval solution (10 mM sodium citrate buffer, pH 6.0) and boiled in microwave oven for 10 min. After cooling down for 30 min, sections were washed twice in dH_2_O. A hydrophobic barrier pen was used to circle the tissue section on the slide before blocking with 5% normal goat serum (NGS) in PBS, for 1 h at room temperature. Subsequently, sections were incubated overnight with primary antibodies diluted in 5% NGS and 0.3% Triton-X in PBS, inside humidified chamber at 4°C. Next day, the sections were washed in PBS three times, with gentle agitation, for 10 min each. Secondary antibody incubation took place inside light-protected jar containing 5% NGS and 1µg/ml DAPI in PBS, for 1h at room temperature. Lastly, sections were washed in PBS, three times for 15 min each, covered with fluorescence-preserving medium and sealed with nail polish. The antibodies used for IF are listed in **Supplementary Table 13**. IF images were acquired with a Leica TCS confocal microscope at 63× magnification.

### Electron microscopy

P24 testes were isolated and fixed in 100 mM sodium cacodylate pH 7.4 with 2% paraformaldehyde and 2.5% glutaraldehyde. Samples were processed and imaged on JEOL JEM-1010 transmission electron microscope, at the Electron Microscopy Resource Laboratory at the University of Pennsylvania.

### Lysate preparation, immunoprecipitation and western blot

P24 testes were homogenized with a pestle in ice cold lysis buffer (20 mM Tris pH 7.5, 200 mM NaCl, 2.5 mM MgCl_2_, 0.5% NP-40, 0.1% Triton X-100 and 1 mM TCEP) supplemented with EDTA-free complete protease inhibitors (Millipore Sigma) and incubated on ice for 5 min. Lysates were further disrupted with sonication 3 times at 30% output (Vibra-Cell, SONICS) and centrifuged at 16,000 g for 15 min at 4°C. Supernatants were flash frozen in liquid nitrogen and stored at −80°C. For IP, lysates were mixed with antibodies against MIWI or MILI and incubated at 4°C for 2 h. Lysis buffer equilibrated Protein G Dynabeads (Invitrogen) were added to the mix and incubated for 90 min at 4°C. Beads were washed 4 times in lysis buffer for 5 min. For WB, protein lysate/eluates were heated in SDS-loading buffer for 12 min at 70°C, resolved on 4-12% NuPAGE Bis–Tris gels (Invitrogen), blotted to nitrocellulose membranes (Invitrogen), blocked with 5% nonfat milk and incubated with primary antibodies overnight. MVH antibody was produced by immunizing rabbits with synthetic peptide SSQAPNPVDDESWD conjugated to KLH protein via an amino-terminal cysteine, followed by affinity purification of sera over columns containing the immobilized peptide (Genscript). All antibodies used for IP and WB are listed in **Supplementary Table 13**.

### piRNA extraction and labeling

The piRNAs bound to immunopurified MILI or MIWI were extracted with TRIzol reagent (Invitrogen) and dephosphorylated by Quick CIP for 10 min at 37°C, in CutSmart buffer (NEB). After CIP inactivation (2 min at 80°C) piRNAs were 5΄end radiolabeled by T4 PNK (NEB) with [γ-^32^P] ATP (10mCi/mL, 3,000 Ci/mmol; Perkin Elmer, BLU502A250UC) in 1× CutSmart buffer supplied with DTT to 5 mM. Labeling reactions were incubated for 30 min at 37°C and stopped with the addition of 2X denaturing gel loading buffer (95% Formamide, 18 mM EDTA, 0.025% SDS, Xylene Cyanol, Bromophenol Blue; ThermoFisher).

### piRNA library preparation

Radiolabeled piRNAs were resolved and gel-purified from 8 M urea 15% polyacrylamide gels (PAGE). Eluted RNA was ligated to miRCat 3′ Linker-1 (IDT) modified with eight extra random nucleotides at the 5′-end. Ligation was performed in 25% PEG 8000 (61) by T4 RNA Ligase 2 Truncated K227Q (NEB) for 8 h at 16°C. Ligated RNA was PAGE purified and reverse-transcribed with a 5′ phosphorylated long primer containing 3′ and 5′ adaptor complementary sequences (62). Primer and adaptor ligated-RNA were heated at 65°C for 10 min and cooled down at room temperature for 5 min. cDNA was produced by AffinityScript (Agilent) in the presence of [α-^32^P] ATP (10mCi/mL, 3,000 Ci/mmol; Perkin Elmer, BLU512H250UC). After PAGE purification, cDNA was circularized by CircLigase I (LGC Biosearch Technologies) and amplified with 25 PCR cycles by Phusion High-Fidelity DNA polymerase (NEB) and adaptor specific primers carrying Illumina p5 and p7 flow cell binding sequences, followed by Illumina 150 Paired-End sequencing in biological replicates. The oligonucleotides used for library construction are listed in **Supplementary Table 14**.

### Ribo-Seq and RNA-Seq

P24 testes were used for preparation of all libraries with the ligation-free method as described by the Sims lab (63), using the SMARTer smRNA kit for Illumina (Takara), with few modifications. For Ribo-Seq, testes lysates were treated with RNAse I and subjected to monosome isolation by S400 spin column centrifugation. Ribosome footprints were purified with 15 % UREA-PAGE. For RNA-Seq, total RNA was used after ribodepletion (Ribominus eukaryotic kit v2; ThermoFisher) and fragmentation (25 min at 94 °C in 2X T4 PNK reaction buffer from NEB –140 mM Tris-HCl pH 7.6, 20 mM MgCl2, 10 mM DTT–) to obtain RNA fragments of similar size to ribosome footprints. RNA fragments (either ribosome protected footprints or RNA-Seq) were *in vitro* polyadenylated and reverse-transcribed with an oligo(dT) primer and a template switching oligo, followed by 12 cycles of PCR amplification with adapter specific primers carrying Illumina p5 and p7 flow cell binding sequences, followed by Illumina 50 Single-End sequencing in biological replicates. Sequencing and library preparation were performed at TB-SEQ.

### RNA extraction, reverse transcription and qPCR

Total RNA was extracted from P24 testes with TRIzol reagent (Invitrogen) and treated with RQ1 DNAse (Promega). 2 µg total RNA was reverse-transcribed by Superscript III with random hexamers (Invitrogen). cDNAs were amplified using PowerUp SYBR-green mix (Applied Biosystems) and the primers listed on **Supplementary Table 15**. Assays were run on StepOnePlus Real-Time PCR system (Applied Biosystems). *Tcp1* was used for input and differentiation stage normalization. Results were calculated from biological triplicates.

### Degradome-Seq library preparation

For Degradome-Seq (PARE-seq), polyadenylated RNA was purified with oligo(dT) beads (NEB) from 15 µg of total RNA extracted from P24 testes. Poly(A) RNA with 5΄phosphates was ligated to a 5’ RNA adapter (CUACACGACGCUCUUCCGAUCUNNN) by T4 RNA ligase I and reverse-transcribed with random hexamers. Libraries were prepared using the NEBNext Ultra II RNA Library Prep Kit (NEB) followed by Illumina 150 Paired-End sequencing in biological replicates. Library preparation and sequencing were performed at CD Genomics.

### Data analysis

The visualizations and statistical analyses were performed in R (v3.6.3). All software tools were installed and run using Conda (v4.11.0) environments. Ensembl genome reference (GRCm38), Ensembl gene annotation, and UCSC RepeatMasker track were used for all analyses. Sequencing batch effects were removed from expression calculation where applicable.

### Quality check, preprocessing, and mapping

Sequencing data quality was checked by FastQC (v0.11.9). The raw reads were preprocessed to remove adapters, low-quality ends, and low-quality reads with Trimmomatic (RNA-Seq, Ribo-Seq, Degradome-Seq) (v0.39) or Cutadapt (piRNA-Seq) (v2.5). Unique molecular identifier sequence (8 nt at the 3’ end) was extracted from piRNA-Seq reads by UMI-tools (v1.0.1), which was also used to deduplicate the mapped reads. Genome alignment was performed with STAR (v2.7.2b). RNA-Seq was mapped with *--alignEndsType Local*. Ribo-Seq and Degradome-Seq was mapped with *-- alignEndsType Extend5pOfRead1*. piRNA-Seq was mapped with *--alignEndsType EndToEnd* and with a maximum of 2 mismatches. Samtools (v1.10), sambamba (v0.7.1), and bedtools (v2.29.0) were used to further process the mapped reads.

### piRNA distribution

piRNA read length was summarized by bbmap (v38.67). Nucleotide composition and genomic origin were determined by piPipes (commit c93bde3). For 3’ end comparisons of piRNAs, mapped MILI-bound and MIWI-bound piRNAs with the same starting genomic position (same 5’ end) with read lengths between 24 nt and 40 nt were used. The median length of each piRNA (per 5’ start position) was determined and the difference in median lengths between *Miwi^+/RK^*and *Miwi^RK/RK^* were calculated and plotted. To calculate 5’-5’ end distances, piRNA reads had to map on the opposite strand within a 30 nt window. The mapped 5’ positions and the distances were determined by bedtools (v2.29.0). Only uniquely mapped reads were considered. The Z-score for 10 nt overlap (ping-pong amplification signature) was calculated with:

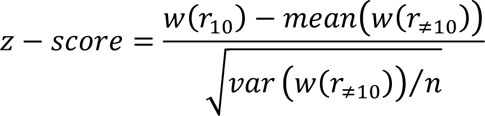

### Transposable elements expression

SquIRE (v0.9.9.92) was used to calculate TEs expression and differential expression. Differential expression was calculated with DESeq2 (v1.16.1) as recommended by SquIRE developers. TEs with adj. p-value <0.05 and fold change >=2 were considered as differentially expressed.

### Gene expression and ribosome occupancy

Transcript expression and ribosome occupancy were estimated with Salmon (v0.14.1). Gene expression was calculated by summarizing transcript-to-gene expression with tximport (v1.12.1) and edgeR (v3.26.0) was used to calculate the differential expression. Differential ribosome occupancy was calculated by edgeR with the formula *∼batch + condition + type + condition:type* formula where: *batch* is the sequencing batch information; *condition* is *Miwi^+/RK^* (Het) or *Miwi^RK/RK^*(RK); *type* is RNA or RIBO; *condition:type* is the interaction between the conditions and types. Genes with adj. p-value <0.05 and fold change >=2 were considered as differentially expressed or with differential ribosome occupancy.

### piRNA expression and targeting

We used the first 25 nucleotides downstream from the first mapped position from the genomic reference as piRNA-representative sequences. piRNA expression was calculated by summing up abundances of all identical piRNA-representatives. edgeR (v3.26.0) was used to calculate the differential expression. piRNAs with adj. p-value <0.05 and fold change >0 were considered as differentially expressed. piRNA targeting was predicted with GTBuster (commit 6697717). The default settings were modified to keep only results, where targeting piRNA is complementary to the targeted transcript with mismatch allowed at the first position, minimum of 14 matches between nucleotides 7-25, and degradome-seq supported cut between 10-11 nt of the piRNA sequence. Only Miwi-expressed transcripts (>=100 RNA-Seq reads in at least one sample) were used to estimate the theoretical targeting. Predicted cuts were overlapped with the UCSC RepeatMasker track by bedtools (v2.29.0). Degradome-Seq supported cuts had to be present in both biological replicates to be considered in subsequent steps. The actual targeted and regulated genes were determined by combing the predicted targeting with gene (adj. p-value <0.1, fold change >0) and piRNA (adj. p-value <0.25, fold change<0) differential expression.

## DATA AVAILABILITY

All datasets have been deposited in the Sequence Read Archive under the BioProject accession PRJNA977257. Link: https://dataview.ncbi.nlm.nih.gov/object/PRJNA977257?reviewer=ac8a7bctmok7hcpth09irvio0v

## CODE AVAILABILITY

Source code utilized for all analyses is available from GitHub at: https://github.com/mourelatos-lab/MIWI-RK_manuscript

## ACKNOWLEDGEMENTS

We are grateful to H. Siomi for 2D9 antibody. This work was supported by National Institutes of Health (NIH) grant GM123512 (Z.M.), Wellcome Trust grant 106144 (D. O’C.), Wellcome Trust Core grant 203149 (Wellcome Centre for Cell Biology), Wellcome Trust multi-user equipment grant 108504 (Wellcome Centre for Cell Biology) and German Research Foundation fellowship DFG ZO 376/1-1 (A. Z.).

## AUTHOR CONTRIBUTIONS

NV, JO, AZ, DO’C and ZM conceived the study. AZ designed, generated and characterized the Miwi-RK mouse with assistance from HY. NV designed, performed and interpreted all other wet lab experiments with contributions by PS, BS, and RF. JO designed, performed and interpreted all computational analyses. All authors analyzed the data. NV, JO, AZ, DO’C and ZM wrote the manuscript with significant input from PS.

## REFERENCES

1. Ozata, D.M., Gainetdinov, I., Zoch, A., O’Carroll, D. and Zamore, P.D. (2019) PIWI-interacting RNAs: small RNAs with big functions. Nat Rev Genet, 20, 89–108.

2. Sarot, E., Payen-Groschene, G., Bucheton, A. and Pelisson, A. (2004) Evidence for a piwi-dependent RNA silencing of the gypsy endogenous retrovirus by the Drosophila melanogaster flamenco gene. Genetics, 166, 1313–1321.

3. Aravin, A.A., Hannon, G.J. and Brennecke, J. (2007) The Piwi-piRNA pathway provides an adaptive defense in the transposon arms race. Science, 318, 761–764.

4. Lewis, S.H., Quarles, K.A., Yang, Y., Tanguy, M., Frezal, L., Smith, S.A., Sharma, P.P., Cordaux, R., Gilbert, C., Giraud, I. et al. (2018) Pan-arthropod analysis reveals somatic piRNAs as an ancestral defence against transposable elements. Nat Ecol Evol, 2, 174–181.

5. Deng, W. and Lin, H. (2002) miwi, a murine homolog of piwi, encodes a cytoplasmic protein essential for spermatogenesis. Dev Cell, 2, 819–830.

6. Kuramochi-Miyagawa, S., Kimura, T., Ijiri, T.W., Isobe, T., Asada, N., Fujita, Y., Ikawa, M., Iwai, N., Okabe, M., Deng, W. et al. (2004) Mili, a mammalian member of piwi family gene, is essential for spermatogenesis. Development, 131, 839–849.

7. Carmell, M.A., Girard, A., van de Kant, H.J., Bourc’his, D., Bestor, T.H., de Rooij, D.G. and Hannon, G.J. (2007) MIWI2 is essential for spermatogenesis and repression of transposons in the mouse male germline. Dev Cell, 12, 503–514.

8. Aravin, A.A., Sachidanandam, R., Girard, A., Fejes-Toth, K. and Hannon, G.J. (2007) Developmentally regulated piRNA clusters implicate MILI in transposon control. Science, 316, 744–747.

9. Reuter, M., Berninger, P., Chuma, S., Shah, H., Hosokawa, M., Funaya, C., Antony, C., Sachidanandam, R. and Pillai, R.S. (2011) Miwi catalysis is required for piRNA amplification-independent LINE1 transposon silencing. Nature, 480, 264–267.

10. Zoch, A., Auchynnikava, T., Berrens, R.V., Kabayama, Y., Schopp, T., Heep, M., Vasiliauskaite, L., Perez-Rico, Y.A., Cook, A.G., Shkumatava, A. et al. (2020) SPOCD1 is an essential executor of piRNA-directed de novo DNA methylation. Nature, 584, 635–639.

11. De Fazio, S., Bartonicek, N., Di Giacomo, M., Abreu-Goodger, C., Sankar, A., Funaya, C., Antony, C., Moreira, P.N., Enright, A.J. and O’Carroll, D. (2011) The endonuclease activity of Mili fuels piRNA amplification that silences LINE1 elements. Nature, 480, 259–263.

12. Vourekas, A., Zheng, Q., Alexiou, P., Maragkakis, M., Kirino, Y., Gregory, B.D. and Mourelatos, Z. (2012) Mili and Miwi target RNA repertoire reveals piRNA biogenesis and function of Miwi in spermiogenesis. Nat Struct Mol Biol, 19, 773–781.

13. Watanabe, T., Cheng, E.C., Zhong, M. and Lin, H. (2015) Retrotransposons and pseudogenes regulate mRNAs and lncRNAs via the piRNA pathway in the germline. Genome Res, 25, 368–380.

14. Zhang, P., Kang, J.Y., Gou, L.T., Wang, J., Xue, Y., Skogerboe, G., Dai, P., Huang, D.W., Chen, R., Fu, X.D. et al. (2015) MIWI and piRNA-mediated cleavage of messenger RNAs in mouse testes. Cell Res, 25, 193–207.

15. Gou, L.T., Dai, P., Yang, J.H., Xue, Y., Hu, Y.P., Zhou, Y., Kang, J.Y., Wang, X., Li, H., Hua, M.M., et al. (2015) Pachytene piRNAs instruct massive mRNA elimination during late spermiogenesis. Cell Res, 25, 266.

16. Dai, P., Wang, X., Gou, L.T., Li, Z.T., Wen, Z., Chen, Z.G., Hua, M.M., Zhong, A., Wang, L., Su, H. et al. (2019) A Translation-Activating Function of MIWI/piRNA during Mouse Spermiogenesis. Cell, 179, 1566–1581 e1516.

17. Hsieh, C.L., Xia, J. and Lin, H. (2020) MIWI prevents aneuploidy during meiosis by cleaving excess satellite RNA. EMBO J, 39, e103614.

18. Wu, P.H., Fu, Y., Cecchini, K., Ozata, D.M., Arif, A., Yu, T., Colpan, C., Gainetdinov, I., Weng, Z. and Zamore, P.D. (2020) The evolutionarily conserved piRNA-producing locus pi6 is required for male mouse fertility. Nat Genet, 52, 728–739.

19. Choi, H., Wang, Z. and Dean, J. (2021) Sperm acrosome overgrowth and infertility in mice lacking chromosome 18 pachytene piRNA. PLoS Genet, 17, e1009485.

20. Vasileva, A., Tiedau, D., Firooznia, A., Muller-Reichert, T. and Jessberger, R. (2009) Tdrd6 is required for spermiogenesis, chromatoid body architecture, and regulation of miRNA expression. Curr Biol, 19, 630–639.

21. Vagin, V.V., Wohlschlegel, J., Qu, J., Jonsson, Z., Huang, X., Chuma, S., Girard, A., Sachidanandam, R., Hannon, G.J. and Aravin, A.A. (2009) Proteomic analysis of murine Piwi proteins reveals a role for arginine methylation in specifying interaction with Tudor family members. Genes Dev, 23, 1749–1762.

22. Kirino, Y., Vourekas, A., Sayed, N., de Lima Alves, F., Thomson, T., Lasko, P., Rappsilber, J., Jongens, T.A. and Mourelatos, Z. (2010) Arginine methylation of Aubergine mediates Tudor binding and germ plasm localization. RNA, 16, 70–78.

23. Tanaka, T., Hosokawa, M., Vagin, V.V., Reuter, M., Hayashi, E., Mochizuki, A.L., Kitamura, K., Yamanaka, H., Kondoh, G., Okawa, K. et al. (2011) Tudor domain containing 7 (Tdrd7) is essential for dynamic ribonucleoprotein (RNP) remodeling of chromatoid bodies during spermatogenesis. Proc Natl Acad Sci U S A, 108, 10579–10584.

24. Meikar, O., Vagin, V.V., Chalmel, F., Sostar, K., Lardenois, A., Hammell, M., Jin, Y., Da Ros, M., Wasik, K.A., Toppari, J., et al. (2014) An atlas of chromatoid body components. RNA, 20, 483–495.

25. Haraguchi, C.M., Mabuchi, T., Hirata, S., Shoda, T., Hoshi, K., Akasaki, K. and Yokota, S. (2005) Chromatoid bodies: aggresome-like characteristics and degradation sites for organelles of spermiogenic cells. J Histochem Cytochem, 53, 455–465.

26. Kotaja, N. and Sassone-Corsi, P. (2007) The chromatoid body: a germ-cell-specific RNA-processing centre. Nat Rev Mol Cell Biol, 8, 85–90.

27. Yokota, S. (2008) Historical survey on chromatoid body research. Acta Histochem Cytochem, 41, 65–82.

28. Hermo, L., Pelletier, R.M., Cyr, D.G. and Smith, C.E. (2010) Surfing the wave, cycle, life history, and genes/proteins expressed by testicular germ cells. Part 2: changes in spermatid organelles associated with development of spermatozoa. Microsc Res Tech, 73, 279–319.

29. Kirino, Y., Kim, N., de Planell-Saguer, M., Khandros, E., Chiorean, S., Klein, P.S., Rigoutsos, I., Jongens, T.A. and Mourelatos, Z. (2009) Arginine methylation of Piwi proteins catalysed by dPRMT5 is required for Ago3 and Aub stability. Nat Cell Biol, 11, 652–658.

30. Nishida, K.M., Okada, T.N., Kawamura, T., Mituyama, T., Kawamura, Y., Inagaki, S., Huang, H., Chen, D., Kodama, T., Siomi, H. et al. (2009) Functional involvement of Tudor and dPRMT5 in the piRNA processing pathway in Drosophila germlines. EMBO J, 28, 3820–3831.

31. Siomi, M.C., Mannen, T. and Siomi, H. (2010) How does the royal family of Tudor rule the PIWI-interacting RNA pathway? Genes Dev, 24, 636–646.

32. Chen, C., Nott, T.J., Jin, J. and Pawson, T. (2011) Deciphering arginine methylation: Tudor tells the tale. Nat Rev Mol Cell Biol, 12, 629–642.

33. Pan, J., Goodheart, M., Chuma, S., Nakatsuji, N., Page, D.C. and Wang, P.J. (2005) RNF17, a component of the mammalian germ cell nuage, is essential for spermiogenesis. Development, 132, 4029–4039.

34. Reuter, M., Chuma, S., Tanaka, T., Franz, T., Stark, A. and Pillai, R.S. (2009) Loss of the Mili-interacting Tudor domain-containing protein-1 activates transposons and alters the Mili-associated small RNA profile. Nat Struct Mol Biol, 16, 639–646.

35. Saxe, J.P., Chen, M., Zhao, H. and Lin, H. (2013) Tdrkh is essential for spermatogenesis and participates in primary piRNA biogenesis in the germline. EMBO J, 32, 1869–1885.

36. Wasik, K.A., Tam, O.H., Knott, S.R., Falciatori, I., Hammell, M., Vagin, V.V. and Hannon, G.J. (2015) RNF17 blocks promiscuous activity of PIWI proteins in mouse testes. Genes Dev, 29, 1403–1415.

37. Ding, D., Liu, J., Midic, U., Wu, Y., Dong, K., Melnick, A., Latham, K.E. and Chen, C. (2018) TDRD5 binds piRNA precursors and selectively enhances pachytene piRNA processing in mice. Nat Commun, 9, 127.

38. Ding, D., Liu, J., Dong, K., Melnick, A.F., Latham, K.E. and Chen, C. (2019) Mitochondrial membrane-based initial separation of MIWI and MILI functions during pachytene piRNA biogenesis. Nucleic Acids Res, 47, 2594–2608.

39. Zhang, H., Liu, K., Izumi, N., Huang, H., Ding, D., Ni, Z., Sidhu, S.S., Chen, C., Tomari, Y. and Min, J. (2017) Structural basis for arginine methylation-independent recognition of PIWIL1 by TDRD2. Proc Natl Acad Sci U S A, 114, 12483–12488.

40. Steger, K. (1999) Transcriptional and translational regulation of gene expression in haploid spermatids. Anat Embryol (Berl), 199, 471–487.

41. Ernst, C., Eling, N., Martinez-Jimenez, C.P., Marioni, J.C. and Odom, D.T. (2019) Staged developmental mapping and X chromosome transcriptional dynamics during mouse spermatogenesis. Nat Commun, 10, 1251.

42. Anzelon, T.A., Chowdhury, S., Hughes, S.M., Xiao, Y., Lander, G.C. and MacRae, I.J. (2021) Structural basis for piRNA targeting. Nature, 597, 285–289.

43. Gainetdinov, I., Vega-Badillo, J., Cecchini, K., Bagci, A., Colpan, C., De, D., Bailey, S., Arif, A., Wu, P.H., MacRae, I.J. et al. (2023) Relaxed targeting rules help PIWI proteins silence transposons. Nature, 619, 394–402.

44. Dowling, M., Homolka, D., Raad, N., Gos, P., Pandey, R.R. and Pillai, R.S. (2023) In vivo PIWI slicing in mouse testes deviates from rules established in vitro. RNA, 29, 308–316.

45. Han, F., Liu, C., Zhang, L., Chen, M., Zhou, Y., Qin, Y., Wang, Y., Chen, M., Duo, S., Cui, X. et al. (2017) Globozoospermia and lack of acrosome formation in GM130-deficient mice. Cell Death Dis, 8, e2532.

46. Eddy, E.M. (2002) Male germ cell gene expression. Recent Prog Horm Res, 57, 103–128.

47. Fanourgakis, G., Lesche, M., Akpinar, M., Dahl, A. and Jessberger, R. (2016) Chromatoid Body Protein TDRD6 Supports Long 3’ UTR Triggered Nonsense Mediated mRNA Decay. PLoS Genet, 12, e1005857.

48. Ellis, P.J., Furlong, R.A., Wilson, A., Morris, S., Carter, D., Oliver, G., Print, C., Burgoyne, P.S., Loveland, K.L. and Affara, N.A. (2004) Modulation of the mouse testis transcriptome during postnatal development and in selected models of male infertility. Mol Hum Reprod, 10, 271–281.

49. Homolka, D., Pandey, R.R., Goriaux, C., Brasset, E., Vaury, C., Sachidanandam, R., Fauvarque, M.O. and Pillai, R.S. (2015) PIWI Slicing and RNA Elements in Precursors Instruct Directional Primary piRNA Biogenesis. Cell Rep, 12, 418–428.

50. Vourekas, A., Alexiou, P., Vrettos, N., Maragkakis, M. and Mourelatos, Z. (2016) Sequence-dependent but not sequence-specific piRNA adhesion traps mRNAs to the germ plasm. Nature, 531, 390–394.

51. Vrettos, N., Maragkakis, M., Alexiou, P., Sgourdou, P., Ibrahim, F., Palmieri, D., Kirino, Y. and Mourelatos, Z. (2021) Modulation of Aub-TDRD interactions elucidates piRNA amplification and germplasm formation. Life Sci Alliance, 4.

52. Boswell, R.E. and Mahowald, A.P. (1985) tudor, a gene required for assembly of the germ plasm in Drosophila melanogaster. Cell, 43, 97–104.

53. Arkov, A.L., Wang, J.Y., Ramos, A. and Lehmann, R. (2006) The role of Tudor domains in germline development and polar granule architecture. Development, 133, 4053–4062.

54. Lasko, P.F. and Ashburner, M. (1988) The product of the Drosophila gene vasa is very similar to eukaryotic initiation factor-4A. Nature, 335, 611–617.

55. Hay, B., Jan, L.Y. and Jan, Y.N. (1988) A protein component of Drosophila polar granules is encoded by vasa and has extensive sequence similarity to ATP-dependent helicases. Cell, 55, 577–587.

56. Kirino, Y., Vourekas, A., Kim, N., de Lima Alves, F., Rappsilber, J., Klein, P.S., Jongens, T.A. and Mourelatos, Z. (2010) Arginine methylation of vasa protein is conserved across phyla. J Biol Chem, 285, 8148–8154.

57. Trcek, T. and Lehmann, R. (2019) Germ granules in Drosophila. Traffic, 20, 650–660.

58. Wei, C., Jing, J., Yan, X., Mann, J.M., Geng, R., Xie, H., Demireva, E.Y., Hess, R.A., Ding, D. and Chen, C. (2023) MIWI N-terminal RG motif promotes efficient pachytene piRNA production and spermatogenesis independent of LINE1 transposon silencing. PLoS Genet, 19, e1011031.

59. Wang, H., Yang, H., Shivalila, C.S., Dawlaty, M.M., Cheng, A.W., Zhang, F. and Jaenisch, R. (2013) One-step generation of mice carrying mutations in multiple genes by CRISPR/Cas-mediated genome engineering. Cell, 153, 910–918.

60. Yang, H., Wang, H., Shivalila, C.S., Cheng, A.W., Shi, L. and Jaenisch, R. (2013) One-step generation of mice carrying reporter and conditional alleles by CRISPR/Cas-mediated genome engineering. Cell, 154, 1370–1379.

61. Munafo, D.B. and Robb, G.B. (2010) Optimization of enzymatic reaction conditions for generating representative pools of cDNA from small RNA. RNA, 16, 2537–2552.

62. Ingolia, N.T., Brar, G.A., Rouskin, S., McGeachy, A.M. and Weissman, J.S. (2013) Genome-wide annotation and quantitation of translation by ribosome profiling. *Curr Protoc Mol Biol*, **Chapter** 4, 4 18 11-14 18 19.

63. Hornstein, N., Torres, D., Das Sharma, S., Tang, G., Canoll, P. and Sims, P.A. (2016) Ligation-free ribosome profiling of cell type-specific translation in the brain. Genome Biol, 17, 149.

